# Novel function of U7 snRNA in the repression of HERV1/LTR12s and lincRNAs in human cells

**DOI:** 10.1101/2023.08.25.554787

**Authors:** Patrycja Plewka, Michał W. Szczesniak, Robert Pasieka, Agata Stepien, Elzbieta Wanowska, Izabela Makalowska, Katarzyna Dorota Raczynska

## Abstract

U7 snRNA is part of U7 snRNP, a complex required for the 3’end processing of replication-dependent histone pre-mRNAs in the S phase of the cell cycle. During this maturation event, the 5’ region of U7 snRNA hybridizes with the highly complementary sequence present in the 3’UTR of histone pre-mRNAs, called histone downstream element, HDE. This base-pair interaction triggers subsequent reactions that eventually result in cleavage and release of mature histone transcripts. Intriguingly, U7 snRNP is constitutively expressed throughout the cell cycle and in nondividing cells, suggesting another function of U7 snRNA/snRNP in cells.

Here, we show that several human endogenous retroviruses (HERVs) are significantly upregulated in HEK293T cells with U7 snRNA knockdown. They predominantly belong to the LTR12 class. Interestingly, some of them are located within long intergenic noncoding RNAs (lincRNAs), which in turn are upregulated in U7 snRNA knockdown cells as well. Significantly, both these HERV1/LTR12s and lincRNAs contain two or more sequence motifs that perfectly match the 5’ end of U7 snRNA, which we called HDE-like motifs. We confirmed that mutations within the HDE-like motifs abrogate U7 snRNA regulatory function and stimulate the expression of selected lincRNAs. Furthermore, we demonstrate that U7 snRNA inhibits HERV1/LTR12 and lincRNA expression at the transcription level. We propose a mechanism in which U7 snRNA hampers binding/activity of NF-Y transcription factor to CCAAT motifs that are frequently found in LTRs as well as in a close proximity to HDE-like motifs.

The expression of many HERV1/LTR12s and lincRNAs regulated by U7 snRNA seems to be tissue specific, therefore, we suggest that U7 snRNA plays a protective role in keeping deleterious genetic elements in silence in selected types of cells.

## INTRODUCTION

U7 snRNA (U7 small nuclear RNA) is synthetized in metazoan cells by RNA polymerase II (RNAP2) and reaches a length of 63 nucleotides in humans. The 3’ region of U7 snRNA forms a stem-loop secondary structure required for its stability, while the central part is occupied by a non-canonical sequence of the Sm-binding site. In the mature U7 snRNP (small nuclear ribonucleoprotein) complex, Sm site is wrapped by Sm/Lsm heptameric protein core which consists of five proteins shared with other U snRNPs (SmB/B’, SmD3, SmE, SmF and SmG), and two unique proteins, Lsm10 and Lsm11 ^1-3^. U7 snRNA and proteins are assembled into the complex in the same maturation pathway as spliceosomal U snRNPs. Thus, after transcription, U7 snRNA precursor is first exported to the cytoplasm, where it is joined with the core proteins and further processed. Next, the particle is reimported into the nucleus ^3-6^.

In contrast to other U snRNPs mostly involved in splicing, U7 snRNP is a key factor involved in the 3’ end processing of replication-dependent histone (RDH) pre-mRNAs ^7^. During this unique maturation event which takes place in the S phase of the cell cycle, RDH pre-mRNAs undergo a single endonucleolytic cleavage after a specific stem-loop structure, located at their 3’ ends and recognized by the stem-loop binding protein (SLBP) ^8,9^. Downstream of the stem-loop structure there is a purine-rich conserved sequence, known as the histone downstream element (HDE), which is highly complementary to the 5’ end of U7 snRNA ^5,10^. The base-pair binding of U7 snRNP to HDE aids in the recruitment of other factors involved in processing, known as the histone cleavage complex. Cleavage occurs between the 3’ stem-loop and the HDE and is catalyzed by the endonuclease CPSF73. Cleavage releases mature histone transcripts, and there is no polyadenylation step involved ^6,11,12^. Moreover, U7 snRNA has already been described as a negative transcriptional regulator. U7 snRNA interacts with transcription factor NF-Y and inhibits its binding to the promoter region of *MDR1* gene in animal cells. The recognition between NF-Y and U7 snRNA was shown to be specific, however the molecular mechanism of this interaction is still unclear ^13^.

Here, we propose another function of U7 snRNA in human cells related to its negative regulation of NF-Y and not linked to the cell cycle. We suggest that U7 snRNA inhibits transcription of specific human endogenous retroviruses (HERVs) and long intergenic noncoding RNAs (lincRNAs), both of which contain U7 snRNA-complementary sequences and CCAAT motifs recognized by NF-Y.

HERVs belong to the ‘retrotransposon’ class of transposable elements (TE) that in total constitute approximately half of the human genome. They resulted from mobile DNA integration events that occurred in the past; however, to date, TEs have mostly lost the ability to transpose. There are ∼500 000 copies of the HERV loci in the human genome and they comprise ∼8% of the genome ^14^. A full-length HERV consists of two long terminal repeats (LTRs) and open reading frames (GAG, POL, and ENV). However, 90% of them exist as solitary LTRs, which nevertheless may contain active regulatory sequences such as promoters, enhancers, splice sites, and polyadenylation signals. Therefore, although transpose-inactive, LTRs can regulate expression of other genes ^14,15^. Moreover, many HERVs are transcribed to form long noncoding RNAs (lncRNAs), and most lncRNAs contain TE sequences, with enrichment of LTR elements ^15-17^. LncRNAs are transcripts longer than 200 nucleotides and almost half of them (∼15 000 genes) belong to lincRNA class with genes located intergenicaly. LincRNAs can be transcribed by RNAP2, spliced, alternatively spliced, capped and polyadenylated. They play various roles in gene expression, both in the nucleus and in the cytoplasm, at the level of epigenetics, transcription, and translation ^18-23^.

HERVs and lincRNAs are generally expressed in a spatio-temporal manner rather than constitutively. As they both act in complex gene regulatory networks, their aberrant level is often associated with human diseases. Therefore, cells need to tightly control their expression. In this study, we show that U7 snRNA mediates in this regulatory pathway through the transcription factor NF-Y. In HEK293T, HeLa and SH-SY5Y cells, U7 snRNA knockdown activates the expression of a specific family of HERV1 (LTR12s) and lincRNAs, suggesting that in those wild-type cells, U7 snRNA plays a protective role in maintaining these genetic elements in a silence state.

## RESULTS

### U7 snRNA regulates the expression of HERV1/LTR12s and lincRNAs in human cells

The function of U7 snRNA is closely connected to the replication phase, where it plays the crucial role in the 3’ end processing of RDH pre-mRNAs. However, the unique components of U7 snRNP complex, U7 snRNA and Lsm11, are expressed throughout the cell cycle (Fig. 1A) ^24,25^. Moreover, U7 snRNA, Lsm10, and Lsm11 are also present in the non-dividing, differentiated SH-SY5Y cells (Fig. 1B) ^26,27^. These suggest that U7 snRNA/snRNP plays other roles in a cell. To address this issue, we performed a knockdown of U7 snRNA in HEK293T cells (U7 KD cells) using an antisense oligonucleotide. The knockdown efficiency reached 90%, as verified by RT-qPCR and Northern blot, and resulted in aberrant 3’ end processing of RDH transcripts (Fig. 1C, Supplementary Fig. S1A-B). After that we analyzed the transcriptomes by high-throughput sequencing (RNA-seq), finding that U7 snRNA depletion resulted in deregulation of hundreds of genes (DEGs, differentially expressed genes), including mostly protein-coding genes and lncRNAs with predominance of lincRNA class (Supplementary Table S1, Fig. 1D). Subsequently, we mapped sequencing reads to genomic regions annotated as repeated sequences. Analysis revealed the differential expression also of these regions (DERs, differentially expressed repeats), with enrichment of HERV1 family (Fig. 1D, Supplementary Table S2). Among DERs and DEGs, all HERVs and almost 90% of lincRNAs were upregulated, suggesting inhibitory effect of U7 snRNA on these genes in wild-type cells. These results were confirmed for randomly selected examples by RT-qPCR (Fig. 1E). The same observations were made in SH-SY5Y and HeLa cell lines with U7 snRNA knockdown, suggesting rather a general phenomenon, regardless of the cell type used (Fig. 1F, G).

**Fig. 1.**
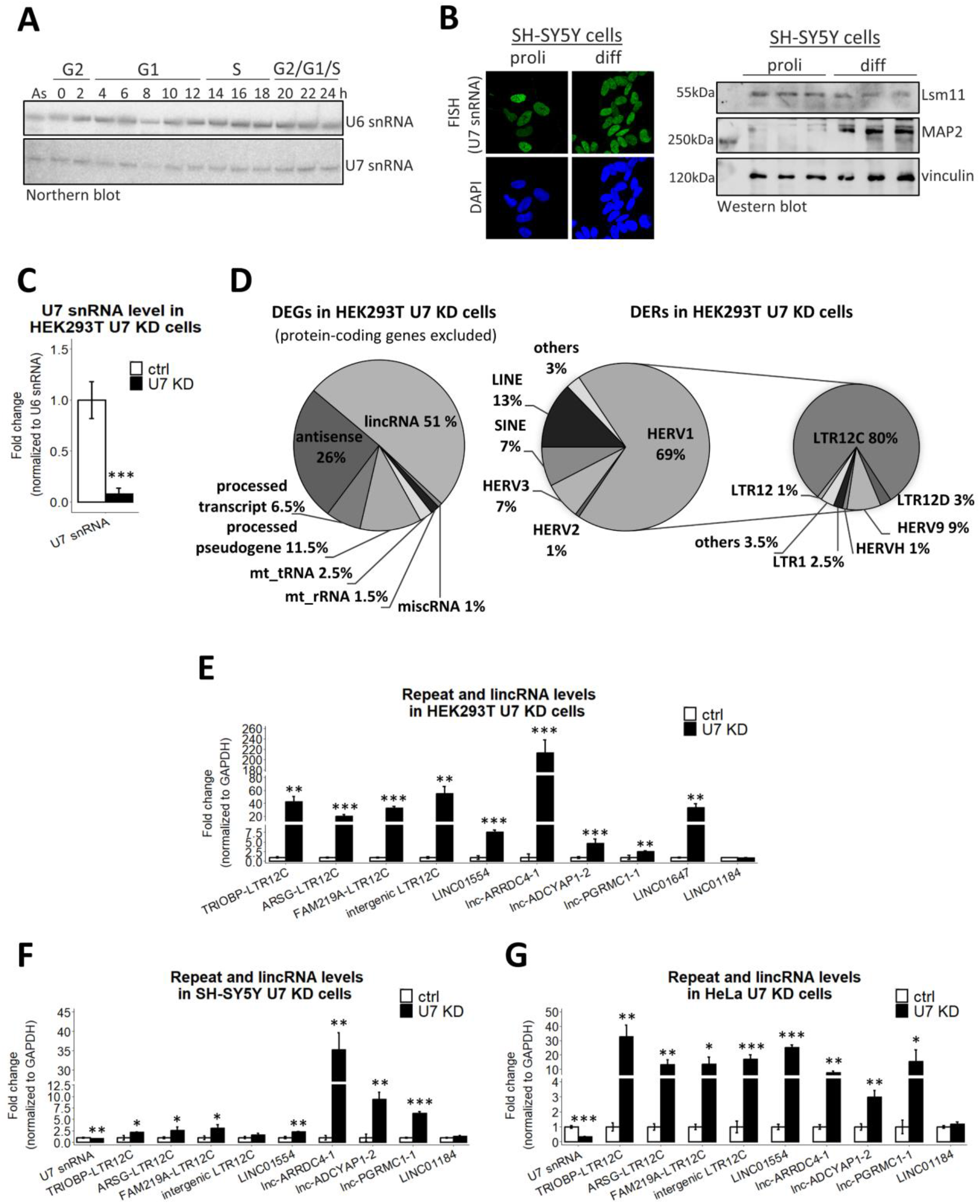
The expression level of U7 snRNA is inversely correlated with the expression of HERV1/LTR12s and lincRNAs. **A)** The level of U7 snRNA at particular phases of the cell cycle was analyzed by Northern blot, U6 snRNA served as a loading control. Synchronization of HeLa cells was performed as described in (^25^, Fig. 3A), cell cycle phases at indicated time points after release from nocodazole block are presented at the top. **B)** U7 snRNA was detected by fluorescence *in situ* hybridization (FISH) and the level of Lsm11 protein was checked by Western blot and immunodetection, in proliferating (proli) and differentiated (diff) SH-SY5Y cells. DAPI was used for nuclear staining. MAP2 was used as a differentiation marker and vinculin was used as a loading control. **C)** U7 snRNA knockdown efficiency in HEK293T cells treated with ASO targeting U7 snRNA (U7 KD) in comparison to control ASO (ctrl) was checked by RT-qPCR. U6 snRNA level served as a normalizer. **D)** Pie charts showing differentially expressed genes (DEGs, on the left) and differentially expressed repeats (DERs, on the right) with special consideration of the HERV1 class, in U7 KD cells. For transparency, protein-coding genes were omitted from DEG analysis. **E, F, and G)** The expression levels of selected HERV1/LTR12s and lincRNAs were analyzed in HEK293T (E), SH-SY5Y (F) and HeLa (G) cell lines, ctrl and U7 KD, by RT-qPCR. LINC01184 was not identified as DEG in RNA-seq data and was used as a negative control. The GAPDH level served as a normalizer. Data represent means ± SD (*n* = 3). P-values were calculated using the Student’s t-test, and the statistical significance is defined as follows: *P ≤ 0.05; **P ≤ 0.01; ***P ≤ 0.001.

Having in mind that lincRNAs can act as molecular sponges (e.g., for miRNAs) ^19,21,22,28^, we examined whether lincRNAs can regulate the accessibility of U7 snRNA throughout the cell cycle stages. We observed that the level of selected lincRNAs is not related to the cell cycle phases; moreover, it remains unchanged in differentiated cells in comparison to proliferating cells (Supplementary Fig. S2A-B). Furthermore, overexpression of selected lincRNAs does not change the level of U7 snRNA nor influence the efficiency of the 3’ end processing of RDH pre-mRNAs (Supplementary Fig. S2C-D), contradicting the hypothesis that the level or accessibility of U7 snRNA is regulated by lincRNAs and pointing at U7 snRNA as the regulator of other gene expression.

### U7 snRNA acts through HDE-like sequence motifs located in HERV1/LTR12s and LTR12-containing lincRNAs

The results mentioned above imply that U7 snRNA may play a role in preserving genome integrity by keeping transposable elements, such as HERVs, silenced. Further analysis revealed that most (84%) of repeated sequences affected by U7 snRNA knockdown (called hereafter U7-dependent, for simplicity) belong to the LTR12 class of HERV1, with over-representation of LTR12C (Fig. 1D, Supplementary Table S2). Since small RNAs operate mainly *via* specific base-pairing with target RNAs, we screened U7-dependent repeats searching for U7 snRNA complementary regions. This resulted in identification of a sequence motif of 15 nucleotides in length exhibiting almost perfect complementarity to the 5’ end of U7 snRNA (Fig. 2A). Due to the similarity to the RDH genes, we called it the HDE-like motif. Further analysis showed that this motif is present within the sequence of all U7-dependent HERV1/LTR12s or in close distance to repeats of DERs from other classes, such as LINEs (long interspersed nuclear elements) or SINEs (short interspersed nuclear elements). Noteworthy, within the human genome, HDE-like motif is over-represented in repeated sequences, with the significant prevalence for LTR12 elements of HERV1 family (81% of all HERVs) (Table 1).

**Fig. 2.**
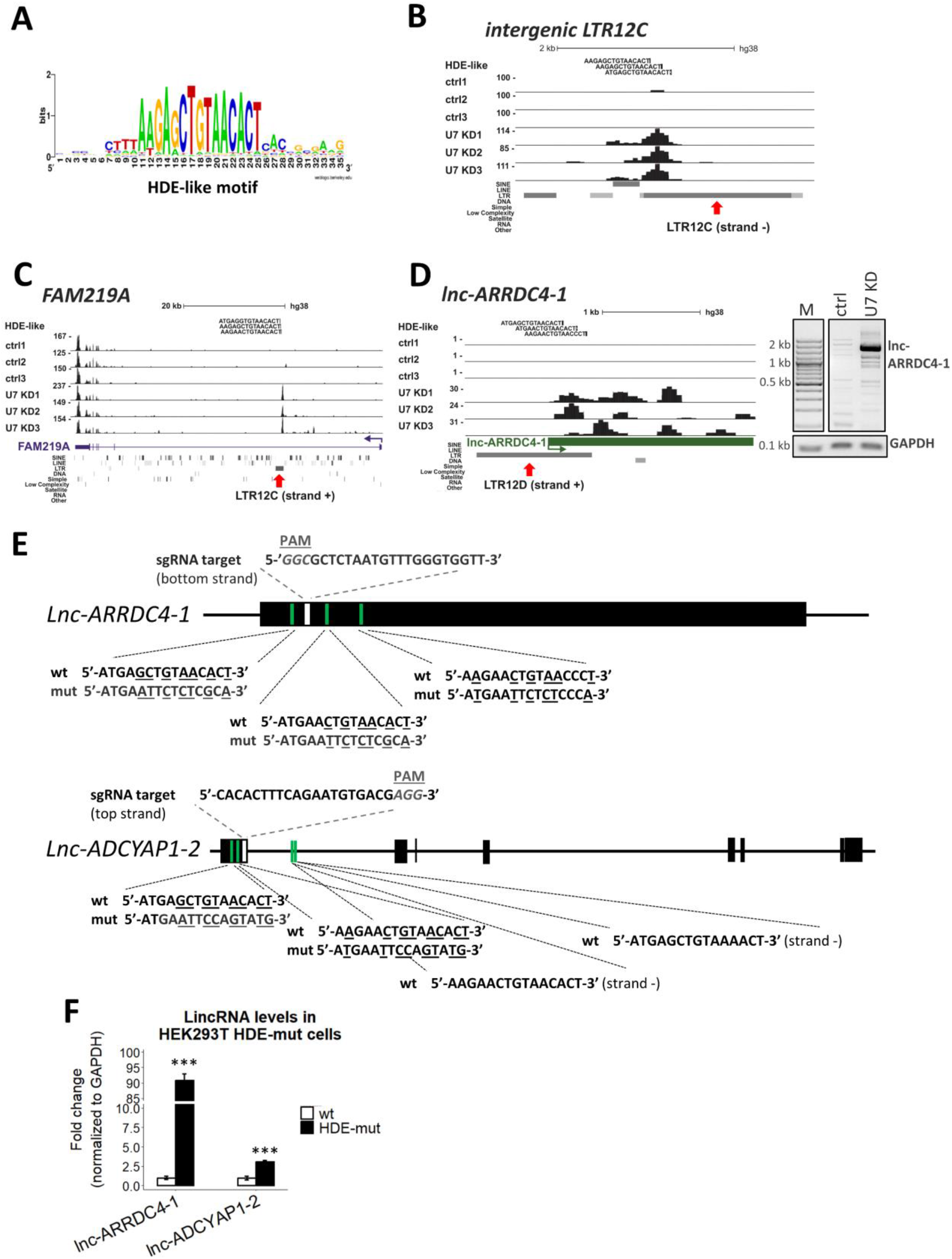
U7 snRNA regulates the expression of HERV1/LTR12s and lincRNAs *via* HDE-like motifs. **A)** Consensus sequence of the HDE-like motif present within HERVs in the context of 10 nucleotides flanking the motif from both sides. **B, C, D)** UCSC Genome Browser visualization of RNA-seq data from HEK293T ctrl and U7 KD cells (three biological replicates are shown) of the intergenic LTR12C element (B), the intragenic LTR12C element in *FAM219A* (C) and the LTR12-containing lincRNA *lnc-ARRDC4-1*; (D). Peaks within data tracks illustrate reads mapped to particular gene regions. Position of HDE-like motifs is shown in the top track. The red arrows indicate the position of the LTR12 elements within the gene structures. The expression of full length lnc-ARRDC4-1 was checked by semi-quantitative PCR, GAPDH was used as a normalizer (D, on the right). **E)** Schematic representation of *lnc-ARRDC4-1* and *lnc-ADCYAP1-2* (isoform ENST00000670380.1). The black boxes represent exons, and the lines depict introns. The positions of HDE-like motifs and sgRNA target sequence used for the generation of HEK293T lnc-ARRDC4-1 HDE-mut and lnc-ADCYAP1-2 HDE-mut cells using the CRISPR-Cas9 system are marked in green and white, respectively. Nucleotides mutated in the HDE-like motifs are underlined. Two HDE-like motifs located within the intron of the *lnc-ADCYAP1-2* gene (in antisense orientation) were not edited. PAM, protospacer-adjacent motif. **F)** The expression levels of the respective lincRNAs in lnc-ARRDC4-1 HDE-mut and lnc-ADCYAP1-2 HDE-mut cell lines were analyzed by RT-qPCR. The GAPDH level served as a normalizer. Data represent means ± SD (*n* = 3). P values were calculated using the Student’s t-test, and the statistical significance is defined as follows: ***P ≤ 0.001.

**Table 1.**
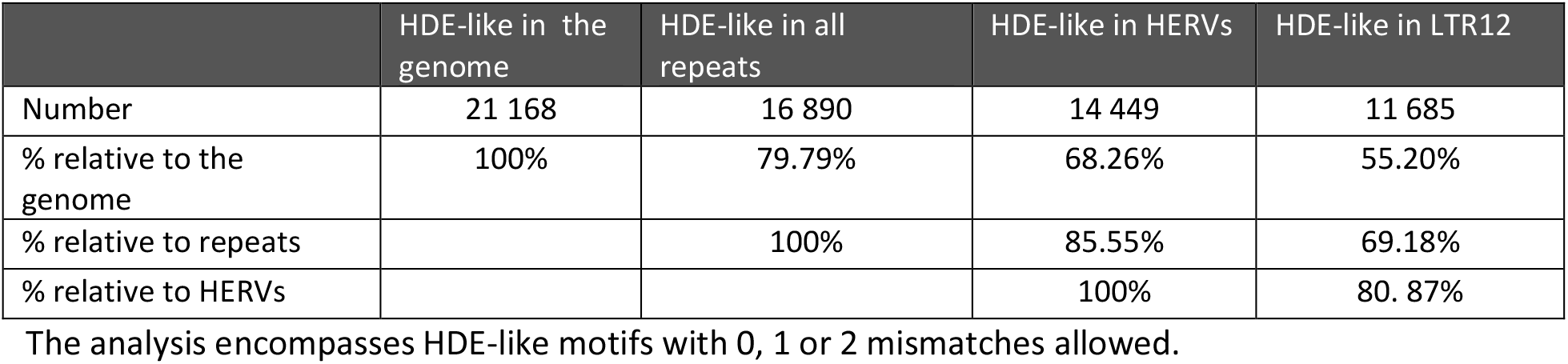
HDE-like motif distribution according to the hg38.

Furthermore, we noticed that U7-dependent repeats are transcribed either as independent molecules (Fig. 2B) or are located within other genes (mainly in introns) (Fig. 2C). In the case of intragenic DERs, the host genes represent mostly protein-coding genes (52%) and lincRNAs (39%). However, by comparing the U7-dependent DERs with U7-dependent DEGs, we noticed that the activation of intragenic-located repeats is rarely accompanied by elevated expression of the host protein-coding gene transcripts (13 per 59). This observation was further confirmed by RT-qPCR showing a specific upregulation of LTR12C elements located in introns of *TRIOBP, ARSG* and *FAM219A* genes in HEK293T cells (Fig. 1E, Fig. 2C), with small or no changes in relative abundance of mature mRNAs and different regions of the host introns (Supplementary Fig. S2E).

As already reported, comprehensive genome-wide studies revealed that TE are present in 75-80% of human lncRNAs, with specific enrichment of HERV/LTR elements. Many of them are important for the function or contribute to the *cis* regulation of lncRNAs ^17,29,30^. Guided by these findings, we analyzed the content of U7-dependent lincRNAs for TE types embedded in them. We found that 50% of U7-dependent lincRNAs contain LTR12 elements with HDE-like motifs. Importantly, in majority of them, HDE-like motifs are present in several copies, predominantly at the beginning of the genes (in the first exons or the first introns) and perfectly or almost perfectly match the 5’ end of U7 snRNA (Table 2). In the case of U7-dependent lincRNAs, either the part containing LTR12 element or the full-length transcript of lincRNA gene is upregulated in U7 KD cells, as exemplified in Fig. 2D. Moreover, distribution profile showing the high accumulation of RNA-seq reads arounds HDE-like motifs within LTR12 elements in U7 KD cells in comparison to wild-type cells (Supplementary Fig. S2F) is in agreement with our previous observation of upregulated expression of HERV1/LTR12s and LTR12-containing lincRNAs in this condition.

**Table 2.**
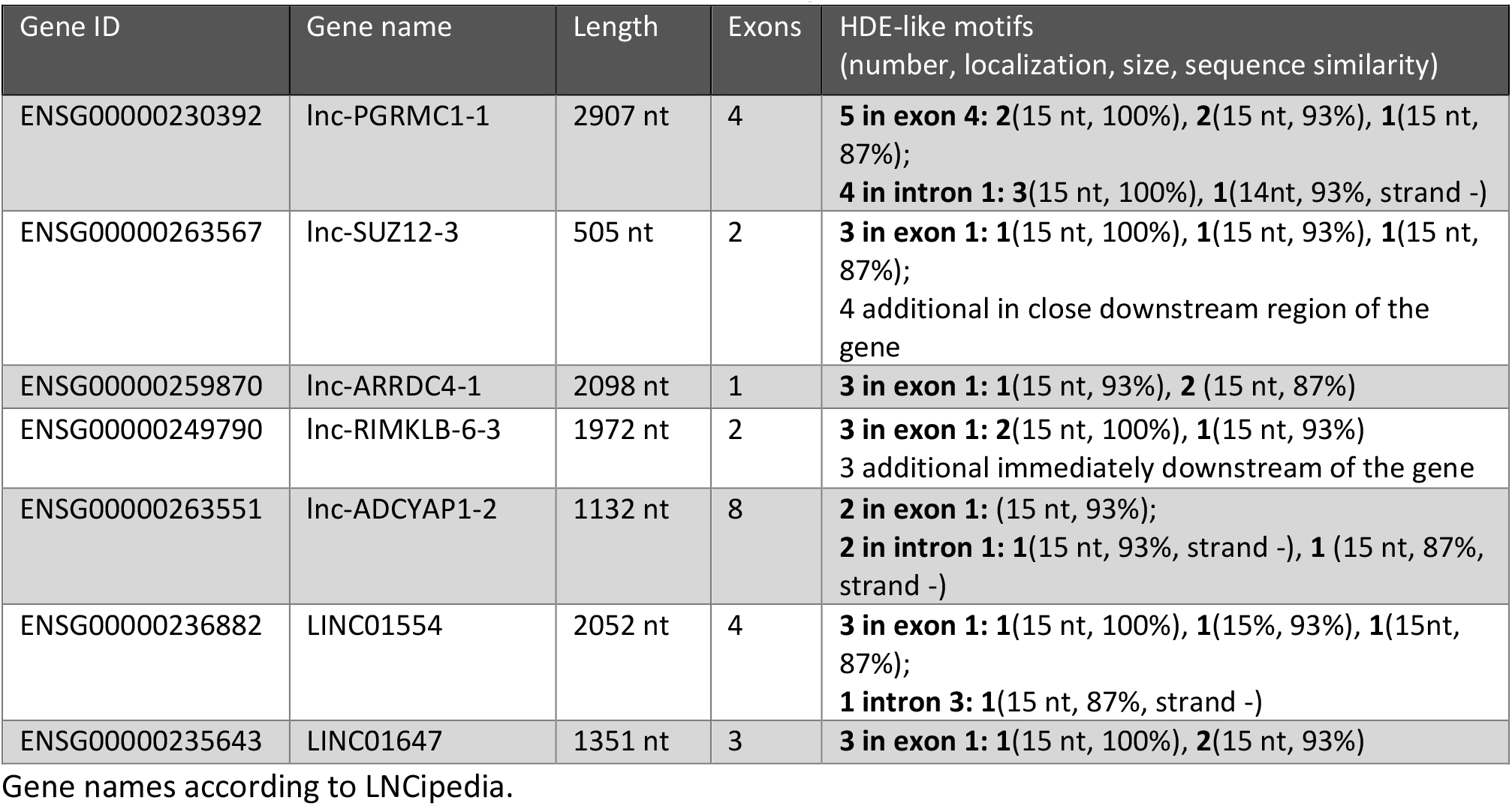
HDE-like motif occurrence in the selected U7-dependent lincRNAs.

Based on the results mentioned above, we suppose that U7 snRNA directly regulates LTR12-containing HERVs and lincRNAs through base-pairing with HDE-like motifs. To test this hypothesis, we used CRIPSR-Cas9 technology and generated cells in which HDE-like motifs were mutated in the *lnc-ARRDC4-1 and lnc-ADCYAP1-2* genes, called hereafter lnc-ARRDC4-1 HDE-mut and lnc-ADCYAP1-2 HDE-mut cells, respectively (Fig. 2E). As shown in Figure 2F, in both HDE-mut cells, the corresponding lincRNAs became almost insensitive to U7 snRNA and are upregulated, confirming a base-pairing interaction between U7 snRNA and HDE-like motifs within the lincRNA sequences.

Next, we addressed the question whether it is U7 snRNA alone or U7 snRNP complex which regulates lincRNA and HERV expression. To answer it, we used cells with inducible knockdown of Lsm10, one of the unique proteins that is the part of U7 snRNP protein core. As shown in Supplementary Fig. S3A, the reduction of Lsm10 mRNA level to ∼40% does not change the level of U7-dependent HERV1/LTR12s and lincRNAs. This result suggests that the mechanism by which U7 snRNA regulates the expression of specific HDE-like containing lincRNAs and HERVs does not require the entire U7 snRNP complex.

### U7 snRNA regulates the transcription of HERV1/LTR12s and LTR12-containing lincRNAs through transcription factor NF-Y

U7 snRNA has been reported to inhibit the activity of NF-Y ^13^. NF-Y is a bifunctional transcription factor capable of activating or repressing transcription and is composed of three subunits, NF-YA, NF-YB and NF-YC, all required for the binding to a CCAAT motif ^31-36^. Interestingly, it has been reported that the HERV1/LTR12 class of transposable elements is abundant in CCAAT motifs, which are located both in tissue-specific enhancers and within the proximal promoter regions ^37^. Likewise, U7-dependent HERV1/LTR12s and LTR12-containing lincRNAs are rich in CCAAT motifs, located usually in the close proximity to HDE-like motifs (Fig. 3A). All these suggest the mechanism in which U7 snRNA regulates the level of HERV1/LTR12s and lincRNAs in cells through the inhibition of their transcription mediated by NF-Y.

**Fig. 3.**
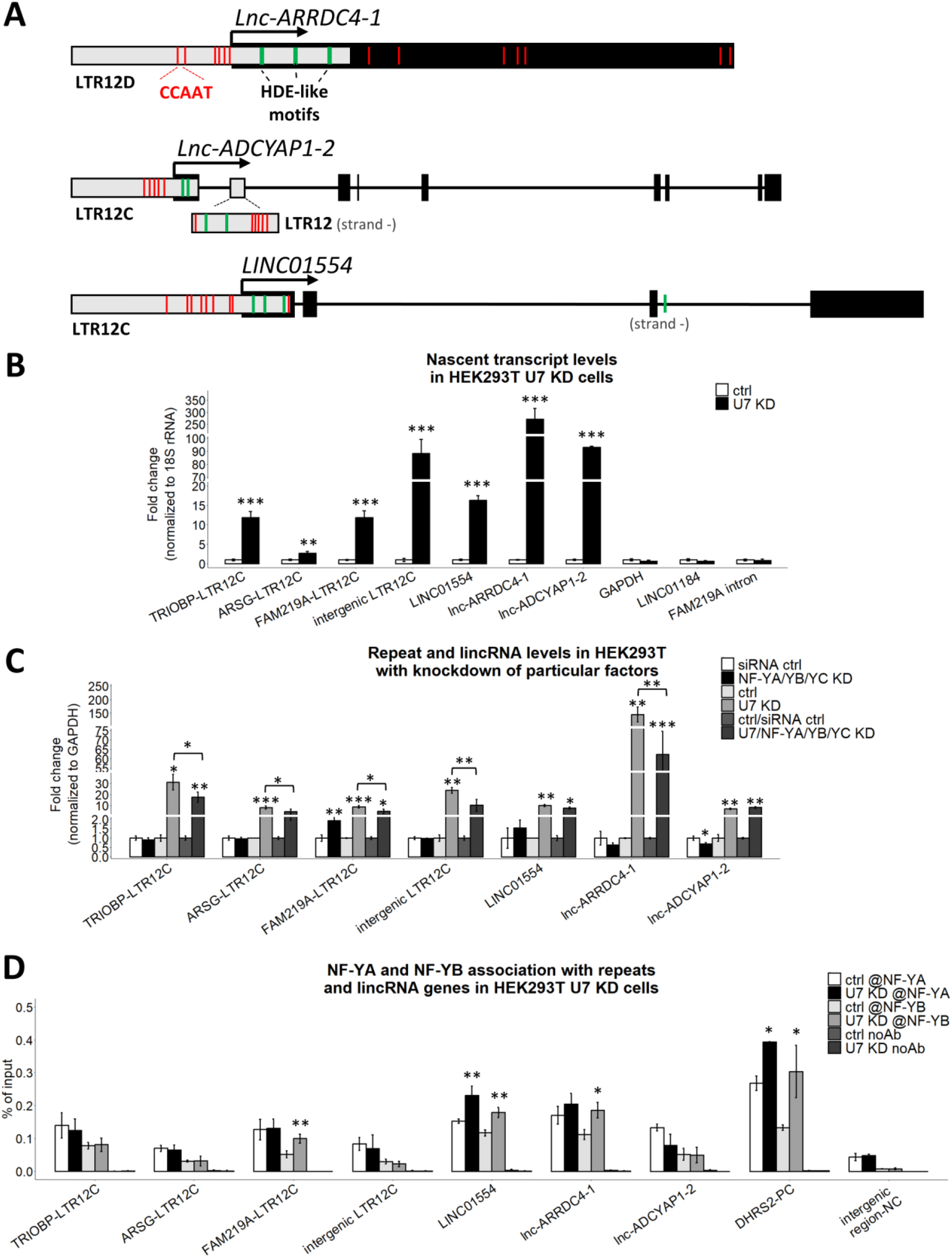
U7 snRNA regulates the transcription of HERV1/LTR12s and LTR12-containing lincRNAs. **A)** Schematic representation of *lnc-ARRDC4-1, lnc-ADCYAP1-2* (isoform ENST00000670380.1) and *LINC01554* (isoform ENST00000436592.5). The black boxes represent exons, the grey boxes illustrate LTR12 elements, and the lines represent introns. The position of the 5’ ends of the genes is indicated by arrows. Positions of the HDE-like and CCAAT motifs are marked in green and red, respectively. **B)** The levels of newly transcribed HERV1/LTR12s and lincRNAs in HEK293T U7 snRNA knockdown cells (U7 KD) compared to ctrl cells (ctrl) were tested by the 4sU assay followed by RT-qPCR. The levels of GAPDH, LINC01184 and FAM219A intron fragment (outside LTR12) were used as negative controls. 18S rRNA was used a normalizer. **C)** The expression levels of selected HERV1/LTR12s and lincRNAs in HEK293T cells with combined depletion of U7 snRNA and NF-Y subunits. The GAPDH level served as a normalizer. **D)** NF-YA and NF-YB binding to HERV1/LTR12s and lincRNA gene regions containing CCAAT sequences in close proximity to HDE-like motifs in HEK293T ctrl and U7 KD cells was analyzed by CHIP followed by qPCR. Empty beads were used as a negative IP control. The bars represent the recovery, expressed as a percentage of the input (the relative amount of immunoprecipitated DNA compared to the input DNA after qPCR analysis). *DHRS2* and an intergenic region were used as a positive (PC) and negative control (NC), respectively. Statistical significance was calculated to assess the association of NF-Y subunits within selected genomic locations between U7 KD and ctrl cells. @-antibody. Data represent means ± SD (*n* = 3). P-values were calculated using the Student’s t-test, and the statistical significance is defined as follows: *P ≤ 0.05; **P ≤ 0.01; ***P ≤ 0.001.

To check whether this is the case, first we measured the transcriptional activity of these genes by 4-thiouridine (4sU) assay. As shown in Fig. 3B, the levels of newly synthesized U7-dependent HERV1/LTR12 RNAs and lincRNAs are significantly higher in U7 KD cells in comparison to control cells. Next, we decided to examine the role of the transcription factor NF-Y in this regulation. First, we knockdown all three subunits of NF-Y (NF-YA/YB/YC KD) in wild-type cells and found no significant alterations in the expression levels of U7-dependent HERV1/LTR12s and lincRNAs. Next, we tested their levels in cells with simultaneous depletion of U7 snRNA and the whole NF-Y complex (Supplementary Fig. S3B) and we figured out that in most cases their synthesis declined in this condition compared to the level observed in U7 KD cells (U7/NF-YA/YB/YC KD vs. U7 KD) (Fig. 3C). This supports the role of U7 snRNA as the inhibitor of NF-Y activity. Then, better understand the role of NF-Y in the repression of U7-dependent genes, we performed chromatin immunoprecipitation (ChIP) using antibodies against NF-Y subunits A and B. Primer pairs were designed to amplify gene regions containing CCAAT sequences in close proximity to HDE-like motifs or intergenic region as a negative control. As shown in Fig. 3D, we observed the binding of NF-YA and NF-YB to HERV1/LTR12s and lincRNAs within regions containing CCAAT motifs. Intriguingly, in some cases, the binding increases in the absence of U7 snRNA, whereas in other cases such obvious changes were not detected although their transcription was increased (Fig. 3B). We suggest that this can be due to partial or full saturation of the given region by NF-Y. When the region is partially saturated by NF-Y, after U7 snRNA knockdown, additional NF-Y subunits can bind to empty motifs and are activated. In turn, when all CCAAT motifs in the given region are occupied, U7 snRNA affects only NF-Y activity. Summarizing, all these results confirm the role of U7 snRNA as an inhibitor of HERV1/LTR12 and lincRNA transcription through the inhibition of binding/activity of transcription factor NF-Y (Fig. 4).

**Fig. 4.**
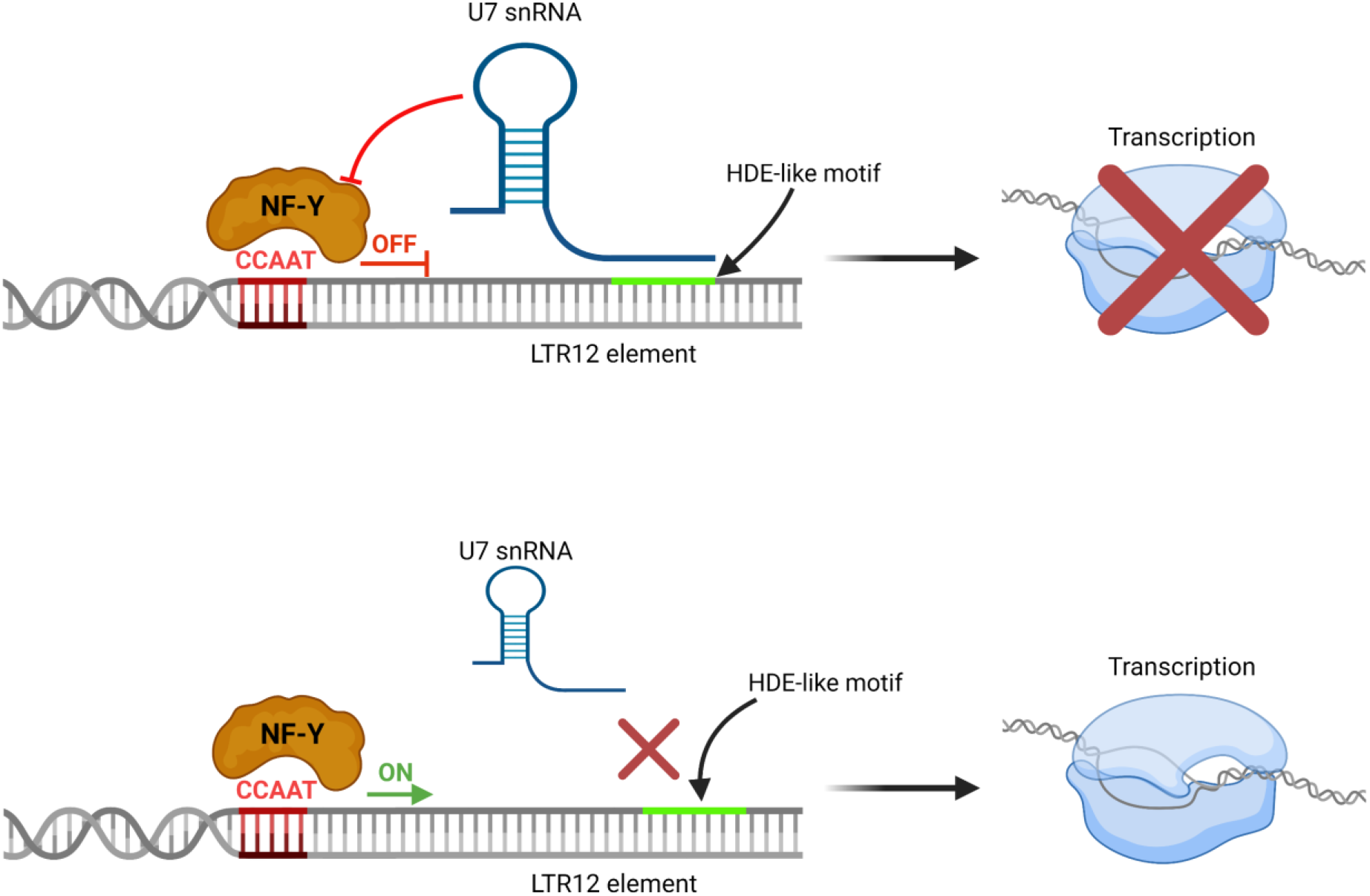
The mechanism of the regulation of HERV1/LTR12 and LTR-containing lincRNA expression by U7 snRNA. In wild-type cells, U7 snRNA base pairs with HDE-like motif present in the LTR12 element and attenuates the binding/activity of transcription factor NF-Y which recognizes CCAAT motif located in the proximity. Transcription of HERV1/LTR12s and LTR12-containing lincRNAs is inhibited (upper panel). In particular conditions (U7 snRNA deficiency, mutations in the HDE-like motif) or cell types, the interaction of U7 snRNA and the HDE-like motif is abolished, NF-Y activates transcription of HERV1/LTR12s and LTR12-containing lincRNAs (lower panel). Created with BioRender.com.

## DISSCUSSION

### U7 snRNA acts as a transcription inhibitor to protect cells from detrimental effect of HERV1/LTR12s and lincRNAs

HERVs and lincRNAs are present in thousands of copies in the human genome, and both are important components of gene expression regulatory networks. In the case of HERVs, this function is mostly related to regulatory sequences present in the LTR regions, and actually most of HERVs exist as solo LTRs. Therefore, although transpose-inactive, HERVs are essential repositories of transcription factor binding sites, such as CCAAT motifs recognized by NF-Y, which may serve as promoters or enhancers. They are also known to produce noncoding RNAs, such as miRNAs, some of which are differentially expressed during human brain development ^14,15,38-41^. In turn, lincRNAs can function both in the nucleus and in the cytoplasm, at the level of DNA, RNA, and proteins. In the nucleus, they can occupy chromatin and be involved in epigenetic regulation and transcription; they can act as protein decoys or scaffolds. In the cytoplasm, lincRNAs modulate mRNA stability and translation. Moreover, a subset of these molecules binds to ribosomes and harbors small open reading frames that may be translated into polypeptides. LincRNAs are also well-known for their function as microRNA sponges ^19-23,42,43^.

As already mentioned, the expression of both HERVs and lincRNAs is tightly regulated. HERVs are mostly silenced in somatic cells including non-dividing, terminally differentiated cells, however, highly expressed in germ cells, during embryogenesis, cell transformation, in neuronal precursors, and pluripotent stem cells (PSC) ^44,45^. Similarly, lincRNAs are expressed primarily in cell type-specific, tissue-specific, developmental stage-specific, or disease state-specific manner, with over-representation in the brain and testis, suggesting involvement in neurogenesis and reproductive functions ^21^. In other kind of cells their expression should be restricted, otherwise they could lead to human diseases. U7-dependent HERV1/LTR12s and lincRNAs also seem to be cell type-specific. Their expression is repressed in some cell lines, such as HEK293T, HeLa, and SH-SY5Y. However, according to genotype-tissue expression (GTEx) portal, many of them exhibit tissue specific expression. This is worth testing in the future as well as elucidating the mechanism by which U7 snRNA can lose its regulatory function in some cells.

An aberrant expression of HERVs associates with chronic inflammatory or degenerative diseases of the nervous system, such as multiple sclerosis, amyotrophic lateral sclerosis (ALS), or schizophrenia as well as with cancer progression ^46,47^. Likewise, mutations of lincRNAs or their deregulation have been linked to tumorigenesis and genetically inherited diseases ^18,28,48^. Unfortunately, U7-dependent lincRNAs and HERV1/LTR12s are poorly studied, and their function remains a mystery. Therefore, it is difficult to speculate what consequences for cells, tissues or even the whole organisms can have their overexpression. Nevertheless, the understanding might shed light on potential therapeutic strategies for neurodegeneration, carcinogenesis, or other human diseases.

Interestingly, the role of one of the U7-dependent lincRNAs, LINC01554, has already been described in hepatocytes, where it exerts tumor suppressive function by regulating cellular glucose metabolism. Its level in hepatocellular carcinoma (HCC) is markedly reduced ^49,50^. Further research revealed that nude mice bearing tumors derived from cells transfected with LINC01554 benefit more from oral administration of the Akt kinase inhibitor MK2206, which is being tested as a therapeutic tool in HCC patients. Therefore, scientists suggest that patients with high expression of LINC01554 could achieve better therapeutic effects from MK2206 ^51^. If this is the case, a similar effect, i.e., an elevated level of LINC01554 in cancer cells might also be achieved by knocking down the U7 snRNA, provided that it plays a regulatory role in this case as well. Noteworthy, LINC01554 is also downregulated in other cancers, including cholangiocarcinoma, testicular germ cell tumors, skin cutaneous melanoma, and ovarian serous cystadenocarcinoma ^51^.

According to our results, U7 snRNA depletion also caused a significant upregulation of two HERV1/LTR12-driven protein-coding genes, *RAE1* and *DHRS2*, which has been known to be positively regulated by NF-Y ^52,53^. Both of these gene promoters contain LTR12 element and also possess HDE-like motifs, showing that not only lincRNAs and HERV1/LTR12 themselves, but also protein-coding gene expression might be controlled by U7 snRNA. In the case of lincRNAs without LTR and HDE-like motifs, we suppose that their upregulation might be an indirect effect of the deregulated multi-layered mechanism involving U7 snRNA action. Anyway, due to their huge functional potential described above, both direct and indirect deregulation of U7-dependent HERV1/LTR12s, lincRNAs, and protein-coding genes might be unfavorable to cells.

### U7 snRNA regulates HERV1/LTR12 and lincRNA expression through a novel mode of action

The novel function of U7 snRNA in human cells proposed here can be played outside of S phase and in addition to its role in RDH genes expression. We showed that U7 snRNA functions as a transcriptional regulator for various genes/regions that possess the CCAAT motif, by the mechanism involving NF-Y, as previously suggested (Fig. 4) ^13^.

However, as shown in Supplementary Fig. S3A, in contrast to its role in the expression of the RDH genes, the mechanism of U7-dependent HERV1/LTR12 and lincRNA expression depends on U7 snRNA alone rather than the canonical U7 snRNP with the Sm/Lsm ring. The observed U7 snRNA reduction in Lsm10 depleted cells is consistent with previously published data ^54^ and is probably caused by decreased stoichiometric requirements for the other components of the U7 snRNP complex due to a lower amount of U7 snRNP particles in general, as a consequence of Lsm10 knockdown. The unassembled U7 snRNA can be degraded by cytoplasmic degradation pathways.

We hypothesize that after synthesis, a pool of U7 snRNA can remain in the nucleus where it hybridizes with HDE-like motifs attenuating NF-Y transcriptional activity. Another pool of U7 snRNA is transported to the cytoplasm and assembled into the U7 snRNP complex. Next, U7 snRNP is re-imported to the nucleus, where together with other proteins from the histone cleavage complex, it participates in the 3’ end processing of RDH gene transcripts. In this process, SLBP stabilizes the base-pair interaction of U7 snRNA with HDE motifs located at the 3’UTR of histone pre-mRNAs which are actually not perfectly complementary to each other. Interestingly, it has been reported that SLBP activity is not necessary for stable interaction in the case of perfect complementarity between U7 snRNA and HDE motif ^55^. In the novel mechanism of regulation described here, most of HDE-like motifs perfectly match to the U7 snRNA (Table 2) and therefore there is no need for additional stabilizing factor. Furthermore, HDE-like motifs are located mainly at the beginning of the lincRNA transcripts, in the first exon or intron, and we demonstrated that U7 snRNA-mediated regulation of these genes and HERV1/LTR12s relies on transcription inhibition rather than on triggering the cleavage, that is next argument confirming novel mode of action playing by U7 snRNA alone, apart from U7 snRNP complex.

The link between U7 snRNA and transposable elements has been described in murine embryonic stem cells (ES) and in preimplantation embryos. The scientists observed that the removal of the 5′ fragment of tRNA-Gly-GCC (tRF-GG) resulted in derepression of ∼50 genes regulated by LTR of the endogenous retroelement MERVL (mouse endogenous retrovirus L) ^56,57^. This was paralleled by decreased synthesis of U7 snRNA/snRNP that in turn led to decreased expression of histones at the transcript and protein level. Although the detailed mechanism was not described, the authors suggest that eventually the lower supply of histones causes chromatin changes and transcriptional activation of the MERVL element, which in turn stimulates the expression of MERVL-linked gene in murine ES cells and preimplantation embryos ^56^. Therefore, this mechanism also differs from the mode of action described here, which is based on the direct regulation of HERV1/LTR12s by U7 snRNA *via* base-pair complementarity with HDE-like motifs and involving transcription factor NF-Y. Such a motif is not present in the MERVL sequence.

### Difficulties with the interpretation of the expression of highly repeated genomic elements

As already mentioned, ∼80% of annotated human lncRNAs have been reported to contain TE sequences, with enrichment of LTR elements ^15,17,29,30^ and we observed similar phenomenon in this report: half of U7-dependent lincRNAs contain LTR12 elements, and ∼40% of intragenic U7-dependent DERs are encoded within lincRNAs. Importantly, some genomic regions that turned out to be affected by U7 snRNA deficiency have their representatives in both U7-dependent lincRNAs and U7-dependent repeats. For example, lnc-PGRMC1-1 (AC004835.1) contains four repeats with the upregulated expression upon U7 snRNA knockdown and has been identified to be activated as the entire gene by DESeq2 analysis as well. It must be pointed out that we cannot rule out the possibility that some lincRNAs that are listed as DEGs, are in fact present on the list only due to the differential expression of the embedded LTR12(s) and *vice versa*. This ambiguity results probably from the imperfection of the algorithm used, which can sometimes identify the gene as DEG based on the sequencing reads mapping only to the one region of the gene, e.g., TE elements. Therefore, it must be further verified in each single case whether the whole lincRNA is upregulated under U7 snRNA knockdown conditions. However, when located at the beginning of the gene, HERV1/LTR12 elements can act as transcriptional activators of this gene, and, indeed, we observed the reads covering the entire gene for the majority of U7-dependent lincRNAs with HERV1/LTR12 elements located in the first exon or the first intron. It is mediated most likely through derepression of the transcription factor NF-Y that has been proposed to play a role in inhibition of HERV1/LTR12 in somatic cells ^37^.

In this report, we also showed that 69% of DERs belong to HERV1 family with LTR12s containing HDE-like motifs, representing the most prevalent class of U7-dependent repeats. In fact, our manual inspection of activated repeats of other types revealed that around 80% of them are located in close vicinity to HDE-like motifs present in neighboring LTR12s. Since DER analysis in U7 KD cells was performed using only uniquely mapped sequencing reads, we assume that a great number of loci activated by U7 snRNA deficiency could have escaped identification due to the high similarity of LTR12 elements in the human genome. Thus, we believe that the novel U7 snRNA-mediated mechanism of HERV1/LTR12 silencing described in this report may have actually a much bigger contribution to cell fitness.

## CONCLUSIONS

We report here that U7 snRNA has a new function in the cell nucleus, in addition to its role in the S phase of the cell cycle played as U7 snRNP complex. Furthermore, we suggest a novel mechanism of regulation of HERV and lincRNA expression in human cells. As summarized in Fig. 4, in this mechanism, U7 snRNA interacts with HDE-likes motifs present within HERV1/LTR12s and LTR12-containing lincRNAs. This, in turn, may attenuate the activity and/or binding of the transcription factor NF-Y to the CCAAT motifs located in the same elements, leading to transcriptional inhibition. Thus, U7 snRNA plays a protective role in keeping these genetic elements in a silent state in specific cell types. However, what phenotypic consequences can be brought about to cells by uncontrolled and pervasive expression of U7-dependent HERV1/LTR12s and specific lincRNAs is still a puzzle due to unknown function of both of them.

## MATERIAL & METHODS

### Cell cultures and transfections

HEK293T, HeLa and SH-SY5Y cells were grown in Dulbecco’s modified Eagle medium with L-glutamine and 4.5 g/L glucose (DMEM; Lonza or Biowest) supplemented with 10% fetal calf serum (Gibco) and antibiotics (100 U/ml penicillin, 100 μg/ml streptomycin, 0.25 μg/ml amphotericin B (Sigma)) at 37°C in a moist atmosphere containing 5% CO_2_. For differentiation into neuron-like cells, SH-SY5Y cells were subjected to 75 μM retinoic acid (USP, Tretinoin, 167400) for 10 days ^58^, with medium replacement every 3 days. The differentiation efficiency was analyzed by measuring the abundance of MAP2 protein (Fig. 1B) or mRNA level of two differentiation markers, RARB and MYC (Supplementary Fig. S2B) ^59^.

For NF-Y knockdown, HEK293T cells were transfected with 20 nM siRNA against NF-YA, NF-YB, NF-YC (Santa Cruz Biotechnology) or 20 nM universal negative control #1 siRNA (Merck), using the Lipofectamine 2000 reagent (Thermo Scientific) according to the manufacturer’s instructions. For U7 snRNA depletion, the chemically modified chimeric ASO ^60^ was introduced to the cells at 100 nM concentration using Lipofectamine 2000, GFP ASO was used as a control (Supplementary Table S3). The cells were harvested and analyzed 48 h post-transfection; knockdown efficiencies were verified by Western blot, RT-qPCR and Northern blot. For overexpression experiments, HEK293T cells were transfected with pcDNA3.1(+)-LINC01554, pcDNA3.1(+)-ARRDC4-1, and pcDNA3.1(+)-ADCYAP1-2 plasmids using Lipofectamine 2000. Cells were harvested 24 h post-transfection. Overexpression efficiencies were verified by RT-qPCR.

HEK293T cells with the mutations in HDE-like elements of *lnc-ADCYAP1-2* (*AP000829*.*1*) (two out of four elements) and *lnc-ARRDCR-1* (*AC024651*.*2*) (three elements) genes were generated according to ^61^ using SpCas9-2A-Puro (PX459) plasmid coding for the respective sgRNAs: 5’-CACACTTTCAGAATGTGACG-3’ or 5’-TTAGTGGGTTTGTAATCTCG-3’; and donor plasmid for HDR (homology-directed repair), and Lipofectamine 2000. 24 h after transfection, puromycin selection at a concentration of 2 μg/ml was applied for 5 days. Clonal cell lines were isolated by serial dilution. Subsequently, colonies grown from single cells were picked, and gDNA was isolated for clone screening using the ExtractMe Genomic DNA Kit (Blirt). The selected clones were genotyped using DreamTaq DNA polymerase (Thermo Scientific) and primers located outside the region spanned by the homology arms to avoid false detection of residual repair template.

HeLa cell line with inducible knockdown of Lsm10 was created using RNAi technology as described in ^62^. Stable cell line expressing inducible scramble shRNA was used as a control. To induce Lsm10 knockdown, cells were treated with 10 μg/ml doxycycline for 48 h.

### RNA extraction, cDNA synthesis, PCR and RT-qPCR

Total RNA was isolated from cells using TRIZOL reagent (0.8 M guanidine thiocyanate, 0.4 M ammonium thiocyanate, 0.1 M sodium acetate pH 5.0, 5% v/v glycerol, 38% v/v saturated acidic phenol) and the Direct-zol RNA MiniPrep Kit (including on-column DNase I treatment, Zymo Research, R2052) or according to the protocol for TRIzol™ Reagent (Invitrogen). For the latter samples, 10 μg of RNA was treated with 2U TURBO DNase (Ambion) for 40-60 min at 37°C followed by standard phenol-chloroform extraction and ethanol precipitation. First-strand cDNA was synthesized in 20 μl reactions with 1-3 μg of RNA using 200 ng of random hexamer primers and 200 U Superscript III reverse transcriptase (Thermo Scientific), according to the manufacturer’s protocol. Semi-quantitative PCR amplifications were performed using DreamTaq DNA Polymerase (Thermo Scientific) and gene-specific oligonucleotide primer pairs. For qPCR, 1 μl of 2-5x diluted cDNA template, 0.2 μM primer mix, and 5 μl of SYBR Green PCR master mix (Applied Biosystems) were mixed in a 10 μl reaction with the following conditions: denaturation for 10 min at 95°C, followed by 40 cycles of 95°C for 15 s and 60°C for 1 min (Applied Biosystems QuantStudio 6 Flex). For testing of U7 snRNA level in U7 KD cells, cDNA was synthesized in a coupled polyadenylation reverse transcription reaction as described in ^25^ (Fig. 1C) or using random hexamers (Supplementary Fig. S1A). Histone pre-mRNA processing efficiency defined as fraction of correctly processed transcript levels was analyzed according to ^25^. The statistical significance of the RT-qPCR results was determined by Student’s t-test. Primers used for PCR and RT-qPCR reactions are listed in Supplementary Table S4.

### Northern blot

For the Northern blot analysis, 30 μg of total RNA were used to detect U7 snRNA. The RNA electrophoresis, blot transfer and hybridization were performed as previously described ^63^. U6 snRNA hybridization signal was taken as a loading control. The sequences of hybridization probes are listed in Supplementary Table S3.

### Fluorescence in situ hybridization

Fluorescence in situ hybridization was performed according to the protocol described in ^59^.

### Metabolic labeling of nascent RNA with 4sU and nascent RNA purification

Metabolic labeling of newly transcribed RNA was performed as described in ^64^ with modifications. Cells were treated with 500 μM 4sU (POL-AURA, PA-03-6085-N#100MG) for 15 min. Nascent RNAs were eluted twice with 100 μl of fresh 0.1 M dithiothreitol (DTT) and precipitated overnight with 2 μl of GlycoBlue (15mg/ml), 20 μl of 3M sodium acetate pH 5.2 and 600 μl of ice-cold ethanol. The precipitated RNA was centrifuged for 30 min at 16 000xg. The pellet was washed twice with 75% ethanol, centrifuged for 10 min at 16 000xg, air-dried and resuspended in RNase-free water. The RNA obtained was reverse transcribed into cDNA as described above. The levels of newly transcribed RNAs were analyzed by RT-qPCR, as described above.

### Western blot and immunodetection

Protein extraction, SDS-polyacrylamide gel electrophoresis (SDS-PAGE) and immunodetection were performed as described in ^65^, using the following antibodies: anti-actin (MP Biomedicals, 691001), anti-NF-YA (Santa Cruz Biotechnology, sc-17753), anti-NF-YB (Santa Cruz Biotechnology, sc-376546), and anti-EWSR1 (Santa Cruz Biotechnology, sc-48404), anti-Lsm11 (Abcam, ab201159), anti-MAP2 (Cell Signaling, 8707), anti-vinculin (Thermo Scientific, MA5-11690).

### Plasmid construction

Plasmid-based donor repair templates for CRISPR were prepared in pGEM-T Easy vector (Promega). Homology arms (each around 800-1000 bp) flanking the site of alterations (HDE-like motifs) were amplified from the genomic DNA using CloneAmp HiFi PCR Premix (Clontech), A-tailed with DreamTaq DNA polymerase and ligated into pGEM-T Easy vector. The mutagenesis of the HDE-like motifs and PAM sequence within donor plasmid for HDR was performed using the QuikChange II Site-Directed Mutagenesis Kit (Agilent Technologies). pcDNA3.1(+)-ARRDC4-1 and pcDNA3.1(+)-ADCYAP1-2 expression vectors were constructed by cloning lnc-ARRDC4-1 and lnc-ADCYAP1-2 sequences to pcDNA3.1(+) vector by using NheI, XbaI and NheI, ApaI restriction sites, respectively. The pcDNA3.1(+) vector containing the LINC01554 sequence was kindly provided by Professor Xin-Yuan Guan ^51^. Primers used for the construction of all vectors are listed in Supplementary Table S4.

### Chromatin immunoprecipitation

Chromatin immunoprecipitation was done as described in ^59^ with some modifications. 225 μl of chromatin (∼5×10^6^ cells equivalent), diluted in a 1:5 ratio using dilution buffer, was used for IP with NF-Y or NF-B antibodies. IP with beads only served as a negative control. One tenth of the chromatin used for immunoprecipitation was used as an input.

### Flow cytometry

Synchronization of HeLa cells in G2/M was done by addition of 200 ng/ml nocodazole (Sigma-Aldrich) to the medium for 18 h. After synchronization cells were washed twice with PBS and then collected 2, 4, 6, 8, 10, 12, 14, 16, 18, 20, 22 and 24 h after release, respectively, as described in ^25^. For G2, G1 and S phase synchronization, HEK293T cells were arrested in G2/M by addition of 200 ng/ml nocodazole and then collected 1.5, 7 and 14 h after release, respectively. For cytofluorometric analysis, the cells were stained with propidium iodide as described in ^62^. Cell cycle profiles were analyzed with a Guava easyCyte System (Merck Millipore) flow cytometer and the data were processed with InCyte Software (utilities from guavaSoft 3.1.1).

### Total RNA-seq library preparation

1 μg of RNA was first subjected to rRNA depletion step using RiboCop rRNA Depletion Kits and next libraries were prepared using Total RNA-Sense Library Preparation Kit (Lexogen), following the manufacturer’s instructions. An Agilent High Sensitivity DNA Kit (Agilent) was used to assess library quality on an Agilent Bioanalyzer 2100, and the libraries were quantified using a Qubit dsDNA HS Assay Kit (Invitrogen). RNA sequencing was performed for 125 bp pair end reads using an Illumina NextSeq HiSeq 2500 platform at the Lexogen NGS facility.

### Bioinformatic analysis

Differential expression of repeats: quality check was done with FastQC v0.11.5 (http://www.bioinformatics.babraham.ac.uk/projects/fastqc/), then reads processing and quality control were done with BBDUK2 v37.02 from BBMAP package (sourceforge.net/projects/bbmap/). Then reads were mapped against the human genome version GRCh38 using STAR v2.5.3a_modified ^66^. Here, due to the nature of repetitive elements, the reads were required to map uniquely (--outFilterMultimapNmax 1 parameter was applied). Also, human genome annotations in a GTF format from ENSEMBL 109 were used to improve the quality of reads mapping. The resulting BAM files were then input to featureCounts v1.5.0-p1 program from Subread software ^67^ to calculate raw expression values of the repeats. Finally, the calculated expression values were subject to differential expression analysis using DESeq2 v1.34.0 from R/Bioconductor ^68^, requiring the adjusted P-value to be below 0.05.

Differential expression of genes: after checking for reads quality with FastQC, they were subject to processing with BBDUK2. Additionally, reads that mapped to human ribosomal RNAs were discarded using Bowtie 2 v2.3.5.1 ^69^. Then, SALMON was used to estimate expression values of genes and transcripts, which was followed by differential expression of genes performed using DESeq2.

Human repetitive elements were downloaded from UCSC Genome Browser via Table Browser utility (track: RepeatMasker, genome: GRCh38, format: BED). The coordinates of HERVs, LTR12s and other repetitive elements were then obtained by parsing the file using scripting languages. HDE-like elements were identified using a custom Python script that was scanning both strands of the human genome (GRCh38), allowing for up to two mismatches compared to the following consensus motif: AAGAGCTGTAACACT.

Analysis of RNA-seq reads coverage around HDE-like elements: the BAM files resulting from uniquely mapped RNA-seq reads were processed with bamCoverage from deepTools v2.5.7 ^70^ to obtain local coverage in vicinity of those HDE-like elements that are located inside LTR12 repetitive elements. The results were then parsed and plotted with a custom Python script using Matplotlib library (https://matplotlib.org/).

## Acknowledgments

We thank Aleksandra Brzek for generating HeLa cells with inducible knockdown of Lsm10, Magdalena Maslon for help in 4sU assay, Kishor Gawade for help with RNA-seq data, and Bartosz Kwiatkowski for help in graph preparation. We also thank Prof. Wojciech Makalowski for stimulating and fruitful discussions about transposable elements.

