## Supplemental Data for "Novel function of U7 snRNA in the repression of HERV1/LTR12s and lincRNAs in human cells"

### Manuscript title

### This PDF file includes:

Supplementary Figures S1 to S3

Supplementary Tables S3 and S4

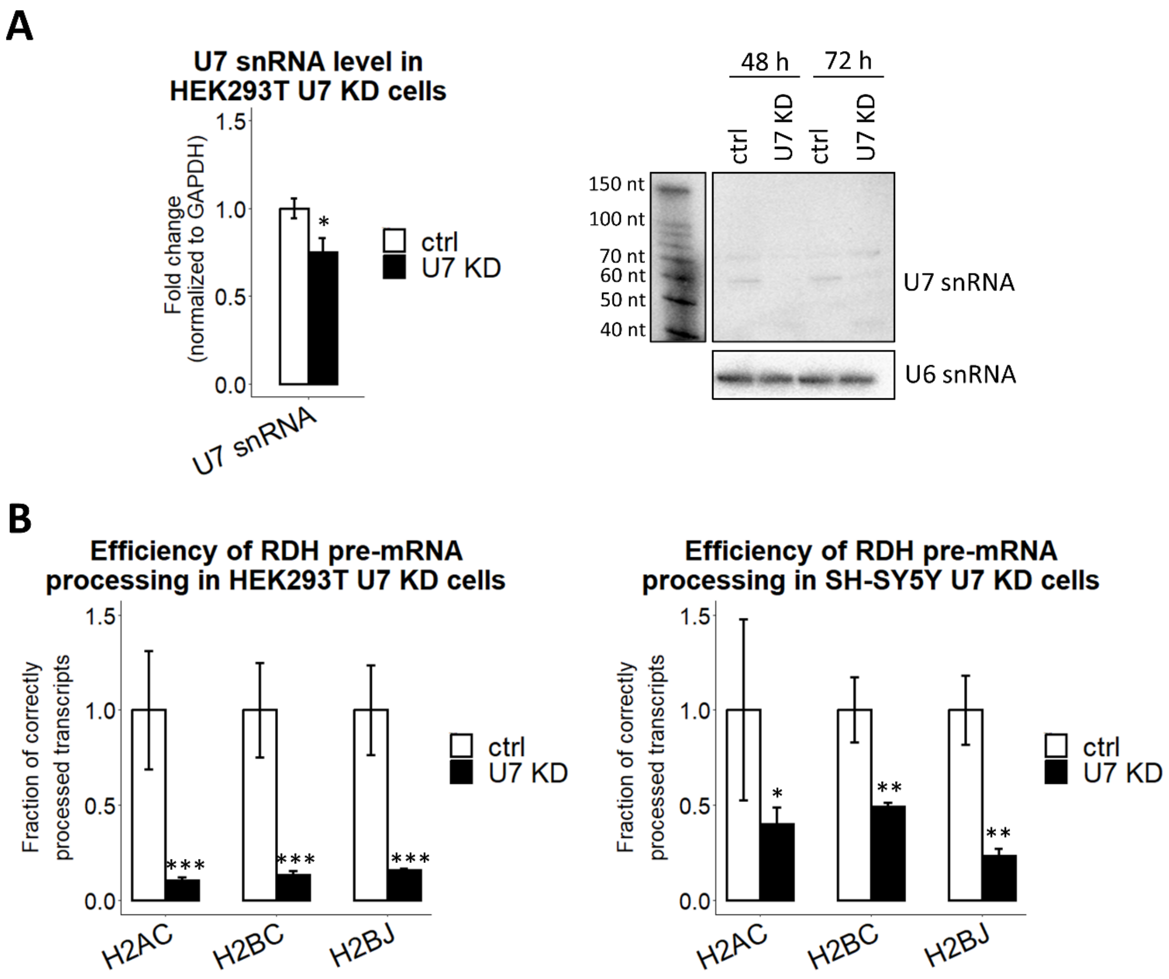


**Supplementary Fig. S1. U7 snRNA depletion results in aberrant 3'-end processing of replication-dependent histone pre-mRNAs. A)** U7 snRNA level in HEK293T cells treated with control ASO (ctrl) or ASO targeting U7 snRNA (U7 KD) was checked by RT-qPCR (with cDNA prepared with hexamer primers) (on the left) and Northern blot (on the right). GAPDH and U6 snRNA levels served as a normalizer and a loading control, respectively. **B)** The fraction of correctly processed histone pre-mRNAs in HEK293T (on the left) and SH-SY5Y (on the right) ctrl and U7 KD cells was assessed by RT-qPCR and normalized to histone H2A.Z pre-mRNA. Data represent means ± SD ($n$ = 3). P-values were calculated using the Student's t-test, and the statistical significance is defined as follows: *P ≤ 0.05; **P ≤ 0.01; ***P ≤ 0.001.

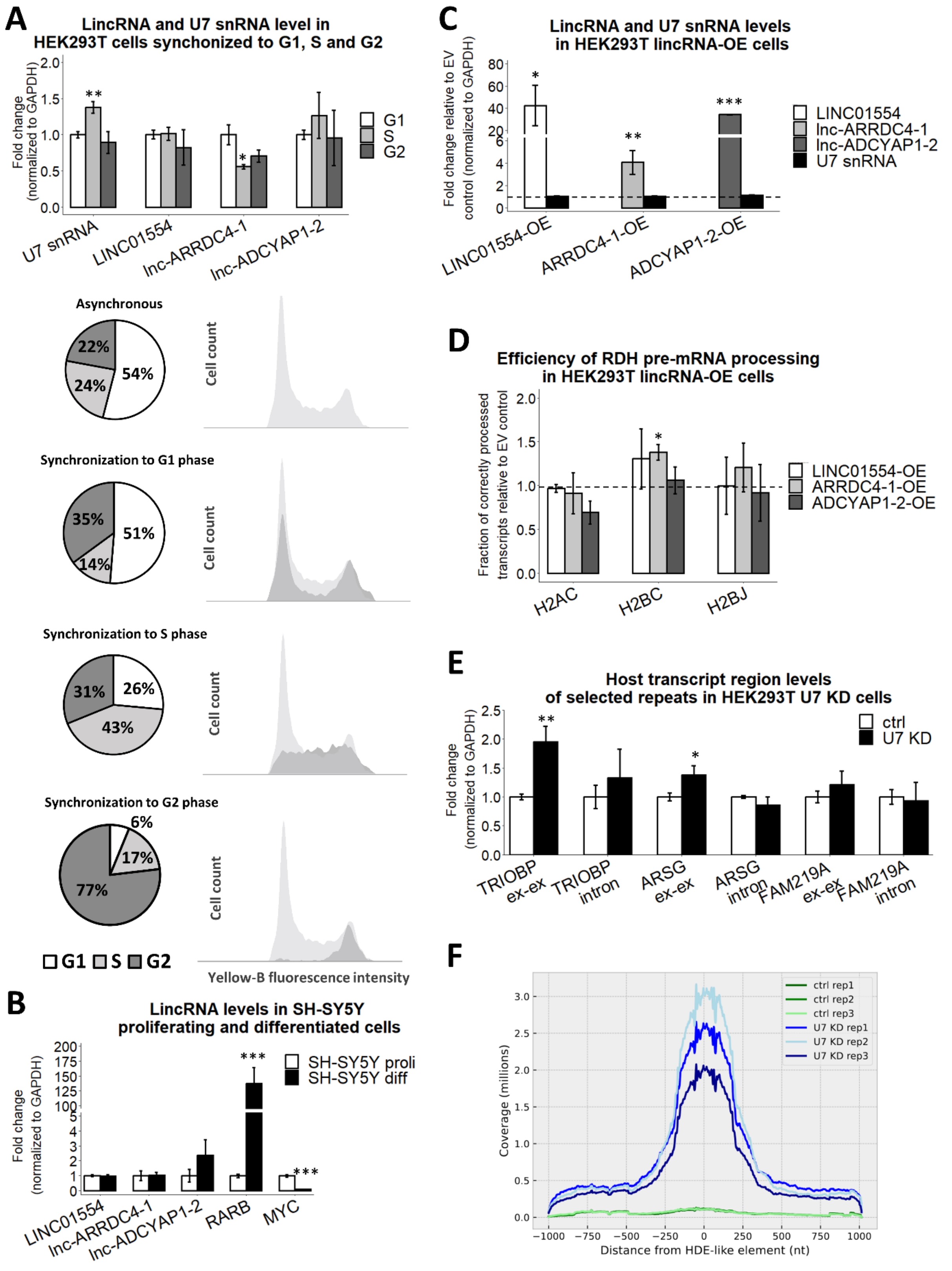


**Supplementary Fig. S2. LINC01554, lncARRDC4-1, and lnc-ADCYAP1-2 do not affect the accessibility of U7 snRNA. A)** LincRNA levels in HEK293T cells synchronized to G1, S, and G2 phases were tested by RT-qPCR (top panel). Flow cytometry analysis of propidium iodide-stained asynchronous and synchronized HEK293T cells. Graphs represent the percentage of cells synchronized to G1, S, or to G2 phase to Yellow-B fluorescence intensity. Light grey color on the histograms illustrates asynchronous cells. **B)** LincRNA levels in SH-SY5Y proliferating (proli) and differentiated cells (diff) were tested by RT-qPCR. RARB and MYC expression served as differentiated and proliferating cell markers, respectively. **C)** U7 snRNA and lincRNA levels in HEK293T cells transfected with pcDNA3.1(+) empty vector (EV) or the particular lincRNA overexpression construct (lincRNA-OE) were tested by RT-qPCR. The GAPDH level served as a normalizer. Dashed line means the level in cells transfected with EV (EV=1). **D)** The fraction of correctly processed histone pre-mRNAs in HEK293T cells transfected with an EV or the particular lincRNA-OE was assessed by RT-qPCR and normalized to histone H2A.Z pre-mRNA. Dashed line means the level in cells transfected with EV (EV=1). **E)** The levels of mRNAs of the host genes or intronic regions in HEK293T cells treated with control ASO (ctrl) or ASO targeting U7 snRNA (U7 KD) were checked by RT-qPCR. The GAPDH level served as a normalizer. Data represent means ± SD (*n* = 3). P-values were calculated using the Student's t-test, and the statistical significance is defined as follows: *P ≤ 0.05; **P ≤ 0.01; ***P ≤ 0.001. **F)** Graph showing the distribution of reads around HDE-like motifs within HERV1/LTR12 elements present in the human genome.

**A**

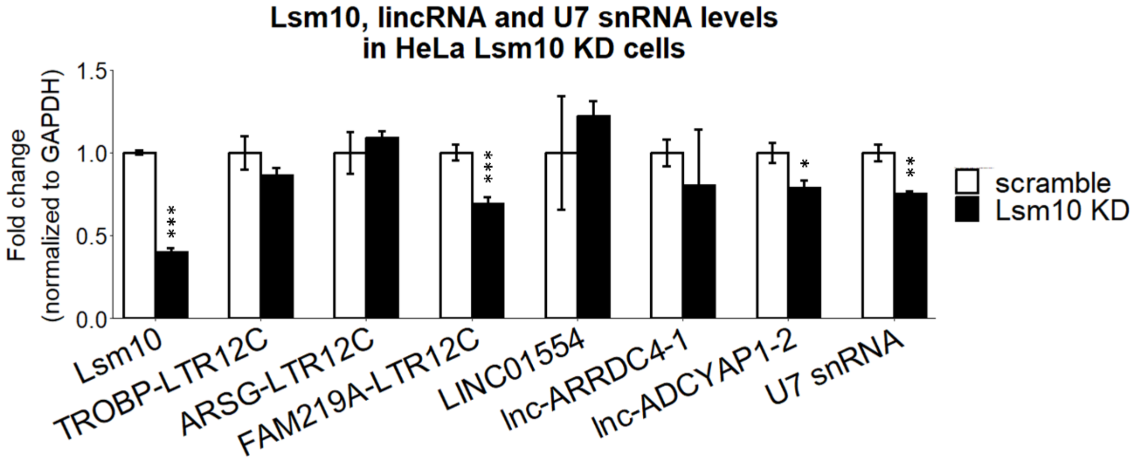


**B**

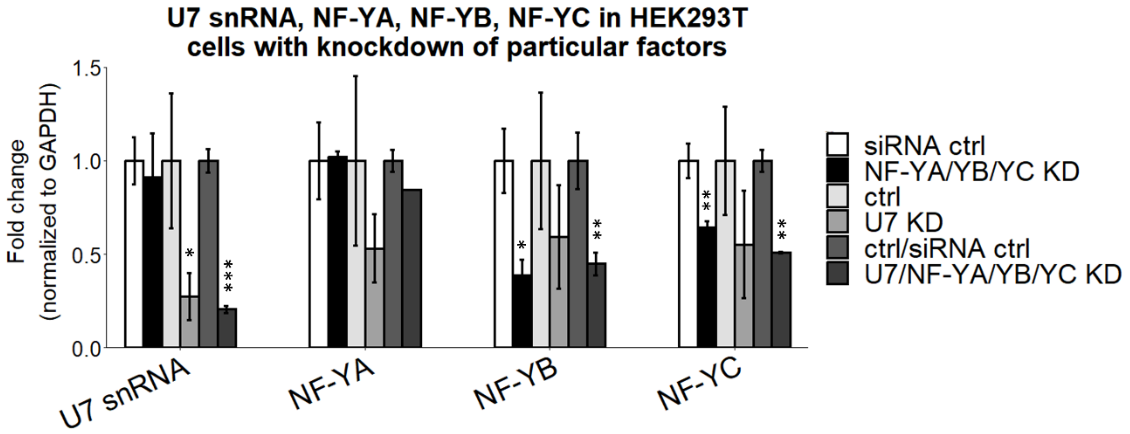


**Supplementary Fig. S3. U7 snRNA, but not the entire U7snRNP complex, is involved in the regulation of HERV1/LTR12s and lincRNAs. A)** Lsm10, HERV1/LTR12, lincRNA and U7 snRNA levels in HeLa scramble and Lsm10 depleted (Lsm10 KD) cells were tested by RT-qPCR. **B)** U7 snRNA, NF-YA, NF-YB and NF-YC levels in HEK293T cells with combined depletion of U7 snRNA and NF-Y subunits. The GAPDH level was used a normalizer. Data represent means ± SD (*n* = 3). P-values were calculated using the Student's t-test, and the statistical significance is defined as follows: *P ≤ 0.05; **P ≤ 0.01; ***P ≤ 0.001.

**Supplementary Table S3. ASOs and Northern blot probes used in this study**

| Name | Sequence (5'-3') |
|---|---|
| **ASO control** | mU*mC*mA*mC*mC*T*T*C*A*C*C*C*T*C*T*mC*mC*mA*mC*mU |
| **ASO U7 snRNA** | mU*mU*mC*mU*mA*A*A*A*G*A*G*C*T*G*T*mA*mA*mC*mA*mC |
| **U7 snRNA_probe** | TTCTAAAAGAGCTGTAACAC |
| **U6 snRNA_probe** | TGGAACGCTTCACGAATTTGCG |

*: phosphorothioate backbone; mN: 2'-O-methylribonucleotide; N: deoxynucleotide.

**Supplementary Table S4. Primers used in this study.**

| Target | Direction | Sequence (5'-3') |
|---|---|---|
| **RT-qPCR** | | |
| TRIOBP-LTR12C | Forward | TAGGGATTTTGCATCCGCCA |
| | Reverse | GCCACCGAAGGTTTTTGAGG |
| ARSG-LTR12C | Forward | GGAAAGTAAGGGGCTAGGTACAG |
| | Reverse | AAGTGGGATCTTGCCAGGTTG |
| FAM219A-LTR12C | Forward | CGAACCCACCAGAAGGAAGA |
| | Reverse | CCTGCAGACGTGTGTCCTAC |
| Intergenic LTR12C | Forward | CGAGGGTCCACGGCTTCAT |
| | Reverse | CTCCAGCCTCGGTGAGAAGG |
| LINC01554 | Forward | TTTCATCTCCAAGAGCGGGC |
| | Reverse | GGCGTTGGAGAGAGGCATAG |
| Lnc-ARRDC4-1 | Forward | ATGCGCCCTAAACACTCCTG |
| | Reverse | ATTGCTCTTCCTTGCGGGAT |
| Lnc-APDCYAP2-1 | Forward | TTCAAAGACTCCAATTTCAGATGGCAAAAT |
| | Reverse | GGTATCATCATAATTGGACCTCTGACTG |
| Lnc-PGRMC1-1 | Forward | GACACTGTACTGGATGTGAGGG |
| | Reverse | GAGCCTTTTTCAAATTTGCACGC |
| LINC01647 | Forward | AGGAAGCATCGGAGACACTC |
| | Reverse | CCCCTGCCTCAGCTCTGTA |
| LINC01184 | Forward | GCGAATCTCCGCTCGTAACT |
| | Reverse | TGAGTCTACCGCTCGAAAGC |
| GAPDH | Forward | CTCAACGACCACTTTGTCAAGCT |
| | Reverse | TCTTACTCCTTGGAGGCCATGT |
| U7 snRNA | Forward | CAGTGTTACAGCTCTTTTAGAATTTG |
| | Reverse | TTCCGGTAAAAAGCCAGAAA |
| DHRS2 | Forward | CACCAAGCGGTGAGACTATCAC |
| | Reverse | CGGGCAACTGCTGACAGCATAG |
| NF-YA | Forward | GGCAGACCATCGTCTATCAACC |
| | Reverse | ATCTGTGCTCCTGCCAAACTGG |
| NF-YB | Forward | GGAATTGGTGGAGCAGTCACAG |
| | Reverse | CCGTCTGTGGTTATTAAGCCAGC |
| NF-YC | Forward | AGTGGCACTGGACAGACCAT |
| | Reverse | CCTGATACAGGCTGGGCTAA |
| HIST2H2AC - total mRNA | Forward | GCAACGACGAGGAACTGAAC |
| | Reverse | GGCTTTGTGGCTTTCGGTTT |
| HIST2H2AC -extended mRNA | Forward | CGAAAGCCACAAAGCCAAAAG |
| | Reverse | GAGCCACCAAAGTGTCAAATG |
| HIST1H2BC - total mRNA | Forward | ATCACCTCCAGGGAGATCCA |
| | Reverse | GAGCCTTTGGGGTTAGGTGT |
| HIST1H2BC -extended mRNA | Forward | TCCAAGTAAGCGTCTTAACACC |
| | Reverse | CCTCTCCAGTTCCTATATTCTA |
| HIST1H2BJ - total mRNA | Forward | CCGAAAAAGGGCTCCAAGAA |

| | | |
|---|---|---|
| | Reverse | CACATAGATGGAATAGCTCTCCTTGC |
| HIST1H2BJ - extended mRNA | Forward | GCTAAGTAAACAGTGAGTTGG |
| | Reverse | CAAGTTACAAGGGTTTGTCAG |
| H2AZ - total mRNA | Forward | GTGTCATTCCACACATCCACA |
| | Reverse | GGCATCCTTTAGACAGTCTTC |
| H2AZ - extended mRNA | Forward | GACATTATTTCCACTCTGGTG |
| | Reverse | GTGCTTAGTTATTGCTGCTAG |
| RARB | Forward | GCGCCTGTGAGGGATGTAAG |
| | Reverse | TGGCATCGATTCCTGGTGAC |
| MYC | Forward | CTGCTTGGACGGACAGGATG |
| | Reverse | GTGAACCAGATCCCGGAGTTG |
| TRIOBP ex-ex | Forward | TGGCCACAAAAGCCTGATCC |
| | Reverse | AGTGTTCACACAGGGCATCC |
| TRIOBP intron | Forward | TGTCCATCCCTGCACCTTTC |
| | Reverse | TGCGACAGCTCTAGAAAGCC |
| ARSG ex-ex | Forward | AGGACACTGCCAACCTTGAT |
| | Reverse | GACTGCAAAGTTGCGTGTGA |
| ARSG intron | Forward | TGAGTTGTGGAAACTTGGACT |
| | Reverse | GGAAATAGGATTTCTGGGTTTCAGG |
| FAM219A ex ex | Forward | GATCGACCGGTTCCAGGAC |
| | Reverse | CTTGGAGCGGGGATGGTTT |
| FAM219A intron | Forward | CTTGCTGGCATTGTTGCAGT |
| | Reverse | GACCCTCCGATAGCCATGTG |
| Lsm10 | Forward | GTAACCACTGTGGACCTGCGG |
| | Reverse | TGCGGCCTGTCACAAAGAGGT |
| **Nascent transcript RT-qPCR analysis** | | |
| LINC01554 | Forward | GTGGGATCACAGAGAGAGCCA |
| | Reverse | ATGTGAAGCAGAGCTGGGTG |
| Lnc-ARRDC4-1 | Forward | ATGCGCCCTAAACACTCCTG |
| | Reverse | ATTGCTCTTCCTTGCGGGAT |
| Lnc-APDCYAP2-1 | Forward | GCGCCGCCTTTTTAAGAACT |
| | Reverse | TCTACATTTCTCTTCCTCATCGTCC |
| 18S rRNA | Forward | GATGGTAGTCGCCGTGCC |
| | Reverse | GCCTGCTGCCTTCCTTGG |
| **CHIP-qPCR** | | |
| LINC01554 | Forward | GTGGGGCCAGATAAGAGAATAAAAG |
| | Reverse | TGGGTTGCCAGTGATGGTTG |
| Lnc-ARRDC4-1 | Forward | CACTCTGTGTCCAGCCAAAC |
| | Reverse | CAAGGCCCCACCACATTAGT |
| Lnc-APDCYAP2-1 | Forward | GCGCCGCCTTTTTAAGAACT |
| | Reverse | TCTACATTTCTCTTCCTCATCGTCC |
| DHRS2 | Forward | GGTCCGTGCCACCTTTAAGA |
| | Reverse | GTGCATTCGTGGTCTTGCTG |
| intergenic region | Forward | TGCTGATAATACTGCTACGAAGGCTG |
| | Reverse | TTTGTGGTTCATCTTTTGAAGTTTCTTTGAGT |

| | | |
|---|---|---|
| **Oligo used in coupled polyadenylation RT reaction and RT-qPCR** | | |
| RT primer | Reverse | GTGCAGGGTCCGAGGTTCAACTATAGGTTTTTTTTTTTTTTTT TT TTTTTTVN |
| universal PCR primer | Reverse | GTGCAGGGTCCGAGGT |
| U7 snRNA | Forward | GCTCTTTTAGAATTTGTCTAGTAG |
| U6 snRNA | Forward | TCGTGAAGCGTTCCATATTTTTAA |
| **Generation of CRISPR/Cas9 mutant cell lines** | | |
| Cloning of sgRNA sequences into SpCas9-2A-Puro vector | | |
| Lnc-ARRDC4-1 | Forward | CACCGTTAGTGGGTTTGTAATCTCG |
| | Reverse | AAACCGAGATTACAAACCCACTAAC |
| Lnc-ADCYAP1-2 | Forward | CACCGCACACTTTCAGAATGTGACG |
| | Reverse | AAACCGTCACATTCTGAAAGTGTGC |
| Cloning of HDR templates into pGEM-T Easy vector | | |
| Lnc-ARRDC4-1 | Forward | CTCTTGAAACGGACGGTTGGA |
| | Reverse | TGTGCATGACAGACACCCTG |
| Lnc-ADCYAP1-2 | Forward | CATTCTGGCCGCACTTGAG |
| | Reverse | GCACTCATTTTCCCTGGACA |
| Mutagenesis of HDR donor plasmids (HDE-like motifs and PAMs) | | |
| Lnc-ARRDC4-1 | Forward | TCCGTGCCACCTTTATGAATTCTCTCGCACACTGCAAAAGTCT A |
| | Reverse | TAGACTTTTGCAGTGTGCGAGAGAATTCATAAAGGTGGCAC GGA |
| | Forward | TCCAGACGCACCACCTTTATGAATTCTCTCGCACACCGTGAA GGTCTGCAGCT |
| | Reverse | AGCTGCAGACCTTCACGGTGTGCGAGAGAATTCATAAAGGT GGTGCGTCTGGA |
| | Forward | ACTGCGGAGACACCATCTTTATGAATTCTCTCCCACACTGCA AGGGGCCGTGGCT |
| | Reverse | AGCCACGGCCCCTTGCAGTGTGGGAGAGAATTCATAAAGAT GGTGTCTCCGCAGT |
| | Forward | TTCATTCCTGAAGCACGCGAGATTACAAAC |
| | Reverse | GTTTGTAATCTCGCGTGCTTCAGGAATGAA |
| Lnc-ADCYAP1-2 | Forward | TGGGTCCACACCGCCTTTATGAATTCCAGTATGCACTGTGAA GGTCTGCAGCT |
| | Reverse | AGCTGCAGACCTTCACAGTGCATACTGGAATTCATAAAGGC GGTGTGGACCCA |
| | Forward | TCTGGACGCGCCGCCTTTTTATGAATTCCAGTATGCACTGTG AGGGTCCGCGGCT |
| | Reverse | AGCCGCGGACCCTCACAGTGCATACTGGAATTCATAAAAAG GCGGCGCGTCCAGA |
| | Forward | CACACTTTCAGAATGTGACGATACCTGAGGTAGAGTTGTGA CT |
| | Reverse | AGTCACAACTCTACCTCAGGTATCGTCACATTCTGAAAGTGT G |
| Genotyping of CRISPR/Cas9 mutant cell lines | | |
| Lnc-ARRDC4-1 | Forward | TGGACAGCACCCGAAACGCTGG |
| | Reverse | GCAAGCATGTGCATACGATGT |
| Lnc-ADCYAP1-2 | Forward | TGGATTGGGAGTTGAAGTGAGT |

| | | |
|---|---|---|
| | Reverse | TGTAACGTCACATCGTTAGATGACT |
| **Construction of lincRNA overexpression plasmids and semi-quantitative PCR** | | |
| LINC01554 | Forward | TCGAGCTAGCCTGTGGAAGCTTTGTTCTTTTG |
| | Reverse | GTCGACTCTAGAGGGGTTCAGGGAACTTCTTAGGG |
| Lnc-ARRDC4-1 | Forward | TCGAGCTAGCGCCCTGTGGAAGCTTTGTTCTT |
| | Reverse | GTCGACGGATCCAAAGCTTCCAGTGTCACCCCCAG |
| Lnc-APDCYAP2-1 | Forward | TCGAGCTAGCGAGTGTGGGAGCTTAGTTCTTTC |
| | Reverse | GTCGACGGGCCCCTGGAGAATGGGAAATCCAAGATC |

N - any nucleotide; V - A or C or G
